# The Critical and Unexpected Role of a Methyl Group in Interleukin-17A Inhibitors

**DOI:** 10.1101/2025.10.02.680113

**Authors:** Xiaobing Deng, Bo Li, Huiling Chen, Gang Zhou, Weiping Lyv, Wenjia Tian, Yaning Su, Yisui Zhou

## Abstract

Interleukin-17 (IL-17) is a pro-inflammatory cytokine primarily secreted by Th17 cells. It plays a crucial role in the body’s immune defense against fungal and bacterial pathogens. However, an imbalance in IL-17 production can contribute to the development of autoimmune and inflammatory disorders. Therapeutic strategies targeting IL-17, such as blocking antibodies like secukinumab (Cosentyx), have been successfully developed. These antibodies are currently employed in the treatment of various conditions, including psoriasis, psoriatic arthritis, and ankylosing spondylitis. More recently, a small molecule inhibitor of IL-17, LY3509754, progressed to clinical trials but was halted during Phase 1 due to unfavorable hepatotoxicity. Two derivatives, compounds 7 and 8, did not advance to clinical trials due to safety concerns. These three compounds (7, 8, and the original lead compound, presumably implied) share a common difluoro substituent, which was hypothesized to be the cause of the observed safety issues. In subsequent structure-activity relationship (SAR) studies, replacing the difluoro substituent with a single methyl group (resulting in compound 9) unexpectedly led to a significant improvement in cellular activity. Furthermore, compound 9 exhibited a very low unbound fraction and reduced liver distribution, ultimately translating to high in vivo efficacy with a sufficient safety margin. This seemingly minor methyl substitution transformed the compound into a highly promising preclinical candidate (compound **9**), now slated for further development. **Co-development** inquiries are welcome. Please contact us at.

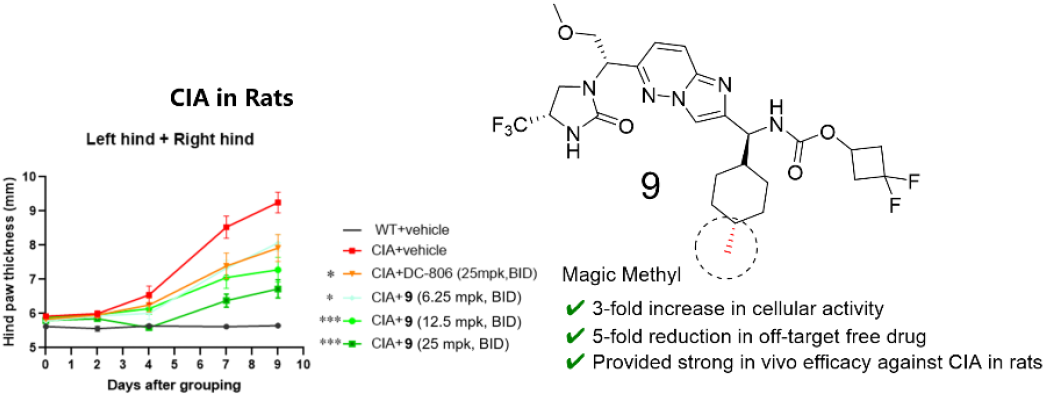

## INTRODUCTION

Interleukin-17A (IL-17A), a pro-inflammatory cytokine primarily produced by T helper 17 (Th17) cells, plays a central role in the pathogenesis of various autoimmune and inflammatory diseases, including psoriasis, psoriatic arthritis, rheumatoid arthritis, and inflammatory bowel disease.^1-3^. As a key orchestrator of immune responses, IL-17-driven inflammation is normally controlled by regulatory T cells and the anti-inflammatory cytokines IL-10, TGFβ and IL-35. However, if dysregulated, IL-17 responses can promote immunopathology in the context of infection or autoimmunity.^4^ Given its critical involvement in these debilitating conditions, the IL-17A signaling pathway has emerged as a validated and highly attractive therapeutic target for immunological disorders.^5^

The clinical success of biologic therapies targeting IL-17A or its receptor has underscored the validity of this therapeutic strategy. Monoclonal antibodies, such as secukinumab (targeting IL-17A), ixekizumab (targeting IL-17A), and brodalumab (targeting the IL-17 receptor A), have demonstrated significant efficacy in treating moderate-to-severe psoriasis and other related inflammatory conditions, revolutionizing patient care.^6-8^ The profound clinical impact of biologic agents targeting IL-17A has validated its therapeutic relevance in human disease, yet the administration via injection remains a significant barrier to patient compliance for a considerable population. This necessitates the exploration of advanced IL-17 therapies, particularly oral formulations, which could offer a more patient-friendly and manageable dosing regimen. Moreover, the pharmacokinetic profile of small molecules presents a distinct advantage over long-acting biologic counterparts. Their comparatively faster systemic clearance, unlike biologics that can exert effects for prolonged durations, allows for a more agile therapeutic strategy. Specifically, the ability to rapidly halt IL-17 antagonism upon the onset of infections, such as those caused by fungi, ensures that the host’s intrinsic immune defenses can be promptly restored, thereby enhancing patient safety. These factors highlight the unmet need for orally bioavailable small molecule inhibitors that can offer greater convenience, potentially lower costs, and broader patient accessibility. The development of small molecule inhibitors targeting the IL-17A pathway presents a unique set of challenges.^9^ Recently, the work of Novartis scientists includes a summary of the discovery process for representative IL-17A small molecule inhibitors **1-6** (Figure 1). ^10^ The compounds **1-3** are too weak to clinical use, and the low solubility of compound **4** limited its clinical exposure. LY3509754 (**6**) had to be discontinued due to concerning drug-induced liver injury (DILI) observed in healthy participants.^11^ Despite the replacement of a potentially hepatotoxic group with an α-fluoroacrylamide moiety in compound **7**, testicular toxicity and hepatotoxicity were still observed, precluding its further development.^10^ This scaffold also attracted the attention of scientists at LEO Pharma, as disclosed in WO2023025783A1, who developed compound **8**. Nevertheless, no clinical advancement for this compound has been documented. Herein, we report our Structure-activity relationships (SAR) study and reveal an unexpected role of a methyl group dramatically altered the potency and pharmacokinetic profile of our compounds. This finding was particularly surprising, as such a minor structural alteration often yields only incremental changes. These findings offer valuable guidance for the future design and optimization of potent and selective small molecule therapeutics targeting the IL-17A pathway.

**Figure 1.**
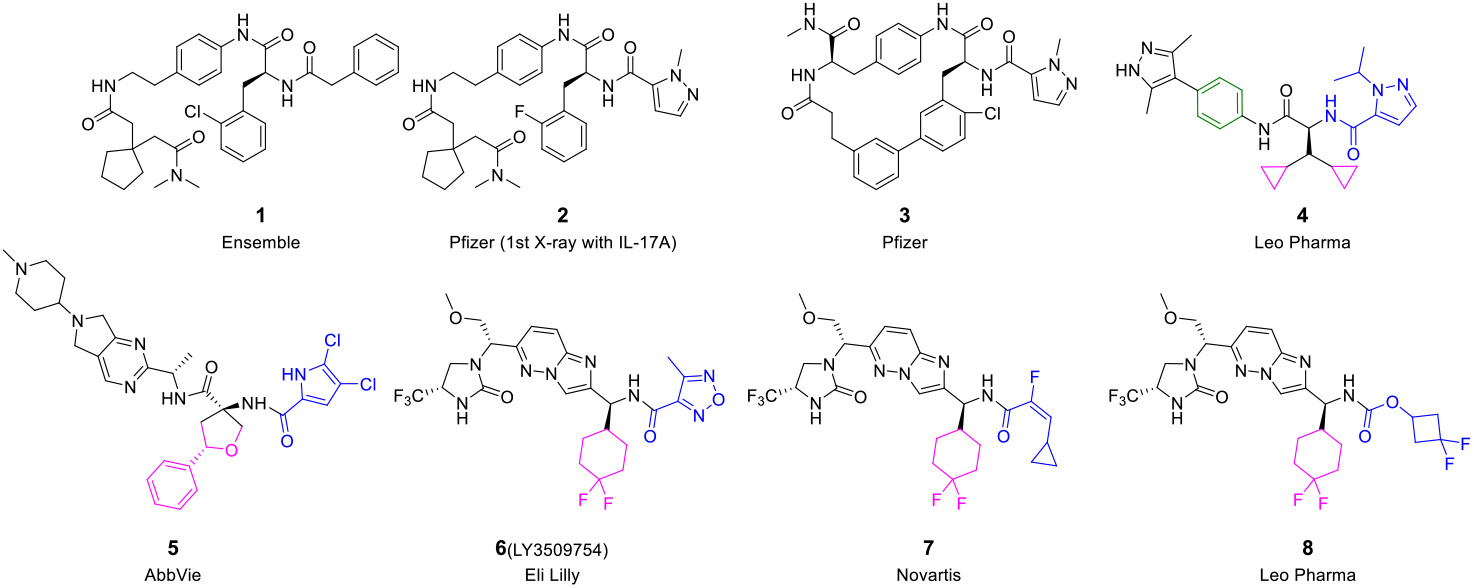
Selected IL-17A low molecular weight inhibitors.

## RESULTS AND DISCUSSION

Both compounds **6** and **7** exhibited hepatotoxicity. Concurrently, LEO Pharma’s compound **8**, which features the same substituent group, did not advance into clinical development. This observation led us to hypothesize that the difluorocyclohexyl moiety might contribute to the observed toxicity. In contrast, DC-806, a known clinical drug candidate, contains a methylcyclohexyl group. Therefore, we embarked on an exploration to replace the difluoro group with a methyl group, aiming to investigate its impact on activity and drug-like properties. Based on publicly available literature and patents, we synthesized the corresponding molecular structures. Our IL-17A/A HEK-Blue assay results demonstrated that compounds **6, 8**, and **10** exhibited similar potency, while compound **9** was nearly three times more potent than these analogues (Table 1). However, IL-17A/RA binding assays revealed that the binding inhibition activity did not entirely correlate with cellular potency. Specifically, DC-806 showed the strongest binding inhibition, while compound **8** exhibited the weakest. This discrepancy likely suggests an allosteric modulation mechanism. Consequently, functional assays might represent a more appropriate evaluation method for IL-17A antagonists, as they better capture the downstream cellular effects rather than just direct binding.

**Table 1.**
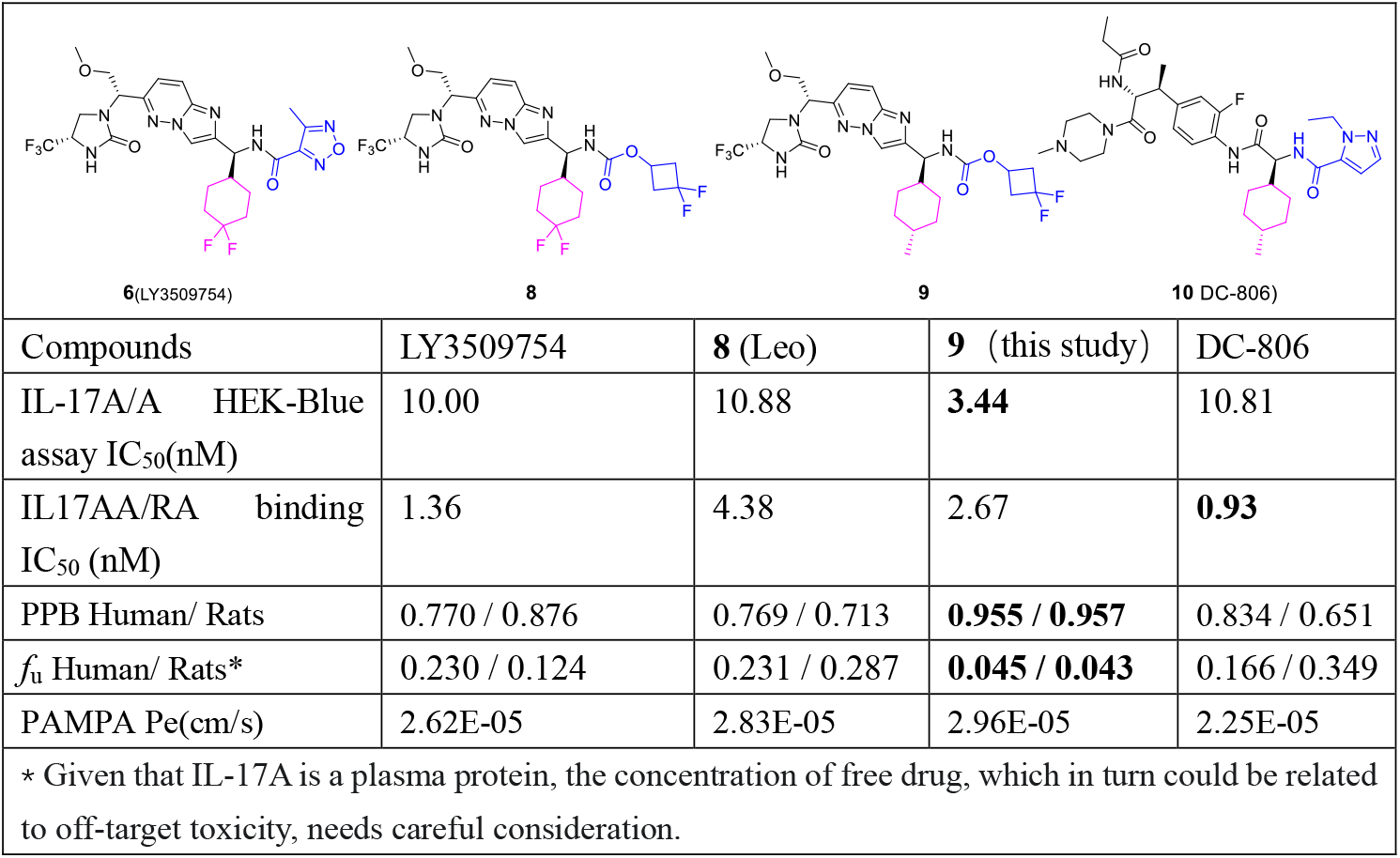
Investigation of Methyl Group Replacement for the Difluoro Group.

|  |  |  |  |  |
| --- | --- | --- | --- | --- |
| Compounds | LY3509754 | **8** (Leo) | **9** (this study) | DC-806 |
| IL-17A/A HEK-Blue assay $IC_{50}$(nM) | 10.00 | 10.88 | **3.44** | 10.81 |
| IL17AA/RA binding $IC_{50}$ (nM) | 1.36 | 4.38 | 2.67 | **0.93** |
| PPB Human/ Rats | 0.770 / 0.876 | 0.769 / 0.713 | **0.955 / 0.957** | 0.834 / 0.651 |
| $f_u$ Human/ Rats* | 0.230 / 0.124 | 0.231 / 0.287 | **0.045 / 0.043** | 0.166 / 0.349 |
| PAMPA Pe(cm/s) | 2.62E-05 | 2.83E-05 | 2.96E-05 | 2.25E-05 |
| * Given that IL-17A is a plasma protein, the concentration of free drug, which in turn could be related to off-target toxicity, needs careful consideration. |  |  |  |  |

Unexpectedly, the methyl-substituted compound **9** exhibited significantly higher plasma protein binding (PPB) compared to other molecules, with plasma protein binding rates of 0.957 and 0.955 in rat and human plasma, respectively. In contrast, other molecules showed PPB rates ranging from 0.65 to 0.87. Consequently, the fraction of unbound drug in plasma (fu) for compound **9** was notably lower at 0.043 and 0.045, while compound **6** displayed higher fu values of 0.124 and 0.23, which likely contributed to its observed hepatotoxicity. As IL-17A is a plasma protein, the concentration of free drug, which in turn could be related to off-target toxicity, needs careful consideration. All four compounds tested were highly permeable, suggesting good oral absorption potential.

Further *in vitro* and *in vivo* metabolic studies were conducted. Rat liver microsome (RLM) stability assays demonstrated that LY3509754 and compound **8** both exhibited metabolic stability. In contrast, DC-806 showed metabolic stability in RLM but exhibited moderate metabolic clearance in human liver microsomes (HLM). Compound **9** demonstrated comparable metabolic clearance to DC-806 in HLM, suggesting that a twice-daily (BID) dosing regimen might be required for future clinical use (Table 2). In the *in vivo* pharmacokinetic (PK) study in rats, all compounds were orally administered at 40 mg/kg. LY3509754 exhibited the highest systemic exposure, with a plasma AUC of 46957 h^·^ng/mL. In contrast, compound **9** and DC-806 showed comparable, much lower plasma AUCs of approximately 5223 h^·^ng/mL. More importantly, regarding the unbound drug AUC (AUC*f*u), compound **9**’s AUC*f*u was merely one twenty-sixth of that of LY3509754. Considering that IL-17A is an extracellular target distributed in the plasma, effectively acting as a plasma protein itself, unbound drug is generally considered unable to effectively target IL-17A. Conversely, unbound drug can reasonably be inferred to be associated with off-target toxicity. Although the mechanisms of drug-induced hepatotoxicity are complex, toxicity is generally dose-dependent. Therefore, reducing drug distribution to the liver might be beneficial in lowering the risk of hepatotoxicity. LY3509754 exhibited a remarkably high liver AUC of 215974 h^·^ng/g. In contrast, compound **9** and DC-806 showed liver AUCs that were less than 30% of LY3509754’s exposure (Table 2). Based on these findings, compound **9** is expected to have a lower risk of hepatotoxicity. Indeed, clinical studies of DC-806 have not reported any instances of liver injury.

**Table 2.** Investigation of Methyl Group Replacement for the Difluoro Group.

| Compounds | Clint<br>RLM / HLM<br>(uL/min/mg) | AUClast (Plasma)<br>Rats 40 mg/kg PO<br>(h·ng/mL) | $AUC_{fu}$ (Plasma)*<br>Rats 40 mg/kg PO<br>(h·ng/mL) | AUClast (Liver)<sup>#</sup><br>Rats 40 mg/kg PO<br>(h·ng/g) |
| --- | --- | --- | --- | --- |
| LY3509754 | 8.53 / < 7.44 | 46957 | 5822.7 | 215974 |
| DC806 | 15.2 / 47.6 | 5416 | 1890 | 64093 |
| **8** (Leo) | 12.9 / < 7.44 | 19538 | 5607.4 | 204463 |
| **9** | 62.1 / 39.4 | 5223 | 224.6 | 61641 |
| * As IL-17A is a plasma protein, the $AUC_{fu}$ (Area Under the Curve of unbound drug) warrants careful consideration, as it may be linked to off-target toxicity; | | | | |
| <sup>#</sup> Liver drug distribution is a critical parameter due to its close correlation with hepatotoxicity. |  |  |  |  |

To comprehensively evaluate the *in vivo* disposition of compound **9**, PK studies were conducted in mice, rats, and monkeys following oral (PO) administration. Due to its exceptionally high intrinsic clearance in dog liver microsomes (Clint: 487 µL/min/mg), the dog species was excluded from further PK studies. The detailed PK parameters are summarized in Table 3.

**Table 3.** Pharmacokinetic Properties of **9** in Different Species.

|  | Mouse |  |  | Rat |  |  | Monkey |  |  |
| --- | --- | --- | --- | --- | --- | --- | --- | --- | --- |
| PO Dose (mg/kg) | 3 | 10 | 30 | 6.25 | 15 | 50 | 3 | 10 | 30 |
| T<sub>1/2</sub> (h) | 1.94 | 1.29 | 2.85 | 2.46 | 2.60 | 1.12 | 7.14 | 4.48 | 3.05 |
| T<sub>max</sub> (h) | 0.44 | 0.61 | 0.83 | 1.0 | 0.83 | 1.0 | 2.0 | 2.0 | 1.67 |
| C<sub>max</sub> (ng/mL) | 343 | 1170 | 3837 | 89.7 | 415 | 2777 | 19.4 | 138 | 618 |
| AUC<sub>inf</sub> (h*ng/mL) | 1062 | 3018 | 18945 | 246 | 1057 | 7268 | 146 | 1017 | 4405 |
| F% | 47.6 | 39.7 | 84.8 | 11.7 | 21.0 | 43.3 | 6.49 | 12.5 | 20.3 |
| Cl<sub>obs</sub> (mL/min/kg)<sup>a</sup> | 23.6 |  |  | 50.1 |  |  | 23.1 |  |  |
| V<sub>ss_obs</sub> (L/kg) | 2.09 |  |  | 2.92 |  |  | 2.98 |  |  |
| * IV doses: mouse, 3 mg/kg; rat, 2 mg/kg; monkey, 1 mg/kg. |  |  |  |  |  |  |  |  |  |

Across all three species, compound **9** exhibited a generally non-linear pharmacokinetic profile following single oral dosing. In mice, doses of 3, 10, and 30 mg/kg yielded AUCinf values of 1062, 3018, and 18945 h^·^ng/mL, respectively. The increase in AUCinf from 10 to 30 mg/kg was disproportionately higher than the dose increase, indicative of non-linear PK at the higher end of the dose range. This non-linearity was further supported by a significant dose-dependent improvement in oral bioavailability (F%), which increased from 47.6% at 3 mg/kg to 84.8% at 30 mg/kg. This trend suggests potential saturation of first-pass metabolism or efflux transporters, leading to enhanced systemic exposure at elevated doses. The observed systemic clearance (CL_obs) in mice was 23.6 mL/min/kg, representing a moderate clearance rate.

A similar non-linear PK behavior was observed in rats. Oral administration at 6.25, 15, and 50 mg/kg resulted in AUCinf values of 246, 1057, and 7268 h^·^ng/mL, respectively. The increase in AUCinf from 15 to 50 mg/kg was notably greater than dose-proportional, confirming the non-linear disposition. Concurrently, the F% improved from 11.7% to 43.3% across the dose range, further supporting saturable processes in absorption or metabolism. The CL_obs in rats was 50.1 mL/min/kg, indicating a moderate to high clearance.

In monkeys, compound **9** also displayed non-linear PK characteristics. Following 3, 10, and 30 mg/kg PO doses, AUCinf values were 146, 1017, and 4405 h^·^ng/mL, respectively. While overall oral bioavailability remained lower than in mice, it also demonstrated a dose-dependent increase, rising from 6.49% to 20.3%, consistent with the non-linear trend observed in other species. The CL_obs in monkeys was 23.1 mL/min/kg, falling within a moderate clearance range.

Across all species, compound **9** exhibited relatively short elimination half-lives (T1/2 ranging from 1.12 to 7.14 h) and rapid absorption, as evidenced by Tmax values typically within 0.44 to 2.0 h. These findings, particularly the consistent moderate clearance rates (23.1–50.1 mL/min/kg) across species, suggest that compound **9** is not subject to excessively rapid elimination, which is generally acceptable for a pre-clinical drug candidate. The observed non-linearity in AUC exposure and F% with increasing doses is a key characteristic of compound **9**’s PK. This phenomenon, likely driven by saturable absorption or first-pass metabolic pathways, implies that target therapeutic exposures could be achieved with higher oral doses, where a greater fraction of the administered dose becomes systemically available. Such properties underscore the importance of careful dose selection for clinical translation and highlight that despite some inter-species variability, compound **9** possesses promising oral absorption potential and acceptable clearance for further development.

The in *vivo* anti-inflammatory efficacy of compound **9** was evaluated in the Collage-induced arthritis (CIA) rat model, with DC-806 serving as a positive control for comparison (Figure 2A). Oral administration of compound **9** demonstrated a clear dose-dependent inhibition of arthritis, as indicated by the progressively increasing efficacy with higher doses. At 6.25 mg/kg BID, compound **9** exhibited 46.20% inhibition of arthritis. This efficacy increased to 72% at 12.5 mg/kg BID and further to 75.20% at 25 mg/kg BID.

**Figure 2.**
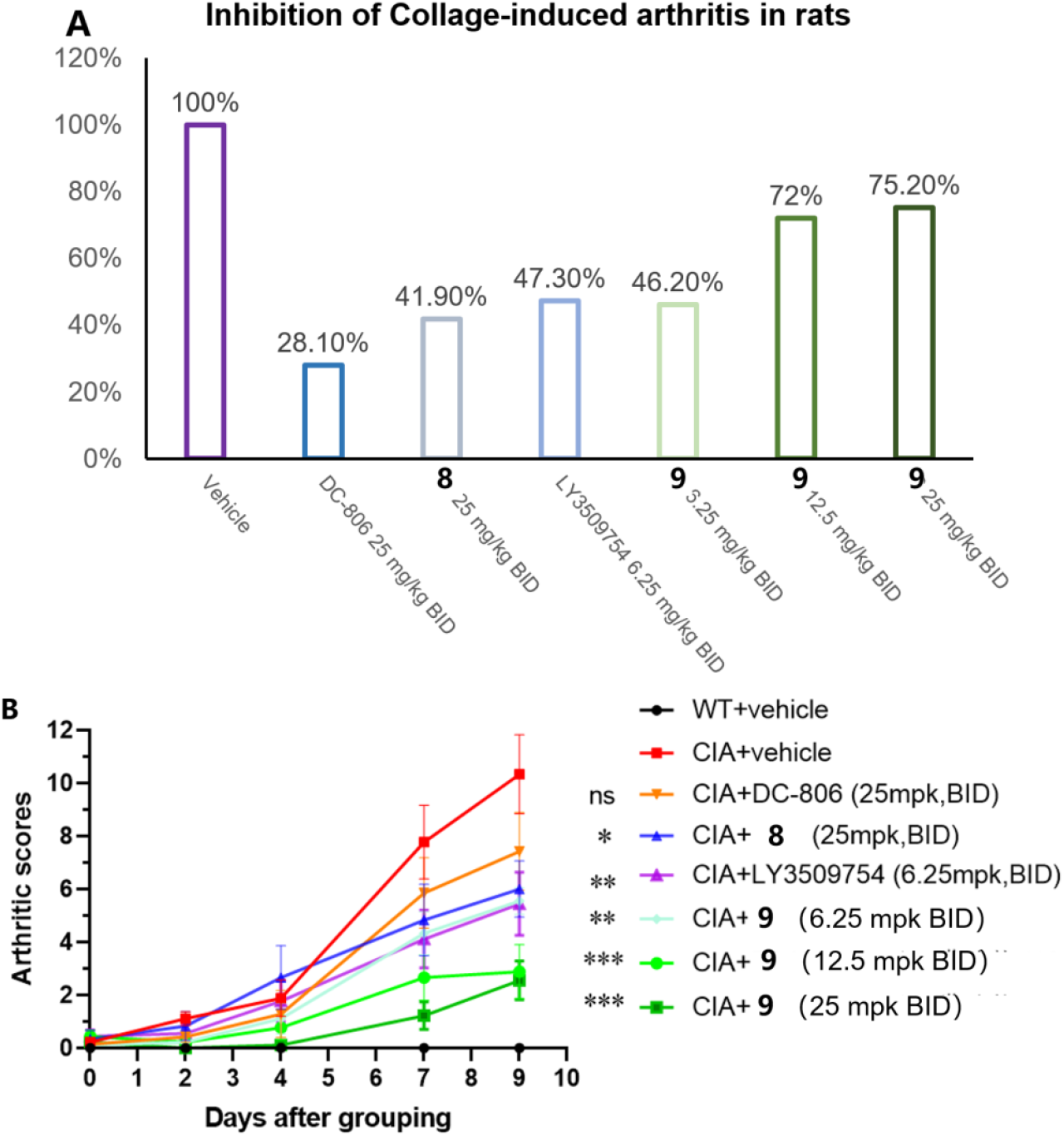
Inhibition of antigen-induced arthritis in rats by selected IL-17A inhibitors.

Notably, even at the lowest tested dose of 6.25 mg/kg BID, compound **9** (46.20% inhibition) showed superior efficacy compared to the positive control DC-806 (28.10% inhibition at 25 mg/kg BID) and compound **8** (41.90% inhibition at 25 mg/kg BID), despite these comparators being dosed at a four-fold higher concentration. Furthermore, the efficacy of compound **9** at 6.25 mg/kg BID was comparable to that of LY3509754 at the same dose of 6.25 mg/kg BID (47.30% inhibition).

This remarkable in vivo efficacy of compound **9**, particularly at lower doses, becomes even more significant when considering its pharmacokinetic profile at the study endpoint (Table 4). Despite achieving comparable or even superior pharmacodynamic effects, the systemic exposure (AUCinf) of compound **9** at 6.25 mg/kg (334 h^·^ng/mL) and 12.5 mg/kg (818 h^·^ng/mL) was substantially lower than that of DC-806 (1869 h^·^ng/mL at 25 mg/kg), compound **8** (2432 h^·^ng/mL at 25 mg/kg), and LY3509754 (7673 h^·^ng/mL at 6.25 mg/kg). For instance, at 6.25 mg/kg, compound **9**’s AUCinf was approximately 23 times lower than LY3509754’s, yet it showed comparable efficacy.

**Table 4.** PK analysis at the study endpoint.

|  | **9** |  |  | DC-806 | **8** | LY3509754 |
| --- | --- | --- | --- | --- | --- | --- |
| PO Dose (mg/kg) | 6.25 | 12.5 | 25 | 25 | 25 | 6.25 |
| T<sub>1/2</sub> (h) | 4.90 | 3.21 | 9.35 | 2.66 | 3.14 | 11.7 |
| T<sub>max</sub> (h) | 0.500 | 0.563 | 0.875 | 0.75 | 0.75 | 0.87 |
| Cmax (ng/mL) | 61.2 | 243 | 1063 | 614 | 613 | 547 |
| AUC inf (h*ng/mL) | **334** | 818 | 4529 | **1869** | **2432** | **7673** |

This striking discrepancy between systemic exposure and potent efficacy for compound **9** can be attributed to its unique properties. Our previous findings indicated that compound **9** exhibits the strongest cellular potency (Table 1) among the tested molecules. Moreover, compound **9** demonstrated a significantly higher plasma protein binding (PPB) rate, resulting in a much lower fraction of unbound drug (*f*u) compared to other molecules. As IL-17A is an extracellular target distributed in the plasma, effectively acting as a plasma protein itself, the bound drug might still contribute to the therapeutic effect or, alternatively, the exceptionally high intrinsic potency of compound **9** allows for strong target engagement even at very low free drug concentrations. These combined characteristics likely underpin the superior *in vivo* efficacy of compound **9** despite its lower systemic exposure in terms of total drug.

The anticipated human dose for compound **9** was projected using allometric scaling of total clearance and volume of distribution at steady-state (Vss) derived from the preclinical PK studies in mice, rats, and monkeys, as detailed in Table 3. Based on these predicted human PK parameters and the potency of compound **9**, which was adjusted relative to the clinically established DC-806, an in vivo effective concentration equivalent to an IC90 of 0.012 μM was targeted. Consequently, the anticipated human dose range for compound **9** was determined to be 70 to 140 mg administered twice-daily.

Following the promising human dose projection, compound **9** was comprehensively evaluated in a battery of early toxicology studies to thoroughly assess its safety profile and potential liabilities. These investigations encompassed a wide range of in vitro and in vivo assays critical for preclinical drug development: GSH conjugation assays, hERG channel inhibition, mini-Ames test, CYP450 inhibition and induction assays. And more advanced hepatotoxicity evaluation was performed in CD34+ human hematopoietic stem cell (huHSC) reconstituted mice, offering a refined in vivo model for assessing liver damage. Finally, a 1-week dose range finding (DRF) toxicology study in rats was conducted at doses of 50 mg/kg and 150 mg/kg to establish initial safety margins and identify potential target organs for toxicity. Compound **9** successfully met all criteria in these rigorous safety assessments, demonstrating a highly favorable toxicological profile. Notably, it exhibited a sufficient and robust safety window, which was significantly superior to that observed for the comparator LY3509754. This extensive preclinical safety evaluation, coupled with its excellent efficacy in the CIA model (Figure 2) and optimized pharmacokinetic properties (Table 3), collectively supported the designation of compound **9** as a preclinical candidate. These comprehensive data underscore its potential for further clinical development as a novel IL-17A inhibitor.

## EXPERIMENTALPART

### Reagents and Materials

HPLC grade methanol and acetonitrile were purchased from Thermo Fisher (Waltham, MA, USA). DMSO, acetic acid, ammonium formate, diclofenac sodium, warfarin, verapamil hydrochloride, α-naphthoflavone, ticlopidine HCl, montelukast sodium, sulfaphenazole, (S)-(+)-N-3-benzylnirvanol, quinidine, and ketoconazole were purchased from Sigma-Aldrich (St. Louis, MO, USA). FaSSIF, FeSSIF, and FaSSGF powders were obtained from Biorelevant (London, UK).

Human liver microsomes and EDTA-K2 plasma were purchased from IPHASE (Beijing, China). NADPH was obtained from Solarbio (Beijing, China). MultiScreen® HTS HV 0.45 µm filter plates, PTFE acceptor plates and MultiScreen-IP PAMPA assay plates were purchased from Millipore (Burlington, MA, USA).

Lecithin was purchased from Yuanye (Shanghai, China), and dodecane was obtained from Alfa Aesar (Haverhill, MA, USA). The dialysis membranes (12-14 kDa) and HTD 96b dialysis device were purchased from HTDialysis (Gales Ferry, CT, USA).

Phosphate buffer (100 mM, pH 7.4) and 50% acetonitrile water was prepared in-house.

The LC-MS/MS system (Sciex Triple Quad 6500) was used for sample analysis. Centrifugation was performed using an Eppendorf 5804R centrifuge (Hamburg, Germany).

MS grade methanol and acetonitrile were purchased from Thermo Fisher (Waltham, MA, USA). DMSO, EDTA-K2 plasma were purchased from IPHASE (Beijing, China).

50% acetonitrile water was prepared in-house.The LC-MS/MS system (Sciex Triple Quad 6500) was used for sample analysis. Centrifugation was performed using an Eppendorf 5910R centrifuge (Hamburg,Germany).

#### Chemistry

All the compounds including LY3509754, DC-806, compound **8** and **9** were synthesized by Beijing double-crane runchuang technology Co. Ltd. 8 was published in WO2023025783A1, and 9 was published in WO2025162215A1.

### IL-17A/A HEK-Blue assay

The IL-17A/AHEK-Blue assay to assess the potential inhibition effect of compounds on IL-17A/A. The Materials, Reagents, Plates and Instrumentation of the Cell Proliferation Assay as shown in the foll owing Table

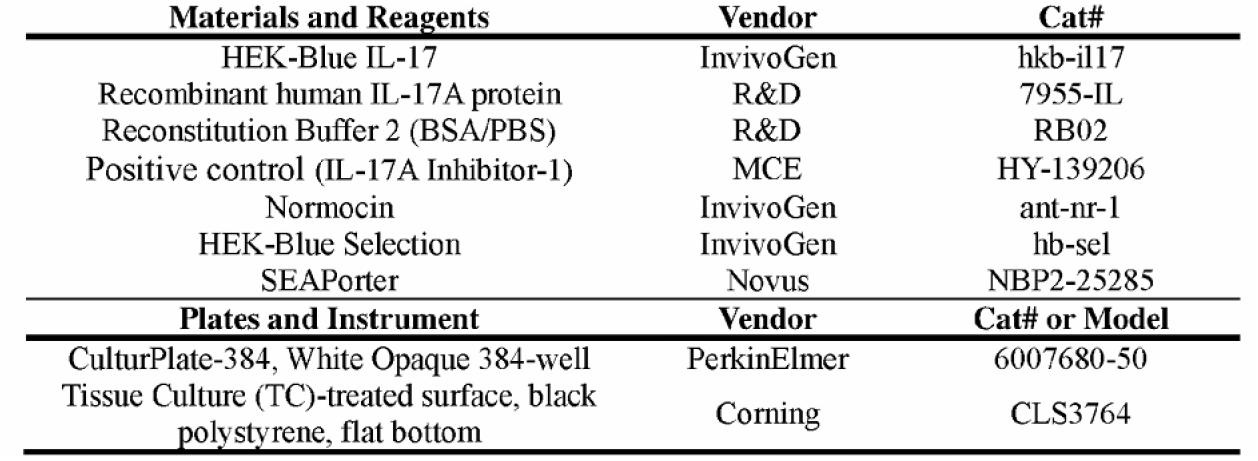

Each test compound was initially solubilized in DMSO to achieve a stock solution concentration of 10 mM. This solution was then subjected to a series of threefold serial dilutions, resulting in a sequence of test solutions of the concentrations at 3.33 mM, 1.11 mM, 0.37 mM, 0.12 mM, 0.04 mM, 0.01 mM, 0.004 mM, 0.002 mM, and 0.5 μM, respectively, for subsequent experimental evaluation. The cultivation of cells was conducted in accordance with the protocols stipulated by the technical data sheet for HEK-Blue™ IL-17, as recommended by the manufacturer. The assay of HEK-Blue IL-17 cells was performed during their exponential phase of growth, adhering to optimal cellular activity conditions. A test solution (25 nl) was dispensed into 384-well plates, followed by the seeding of 25 μl of HEK-Blue IL-17 cells with rhIL-17 protein (10 ng/mL). The resulting cell density in each well was 15,000 cells/well. cells were incubated for 20 hours at 37°C under a 5%CO2 atmosphere. The cell supernatant (2 μL) was transferred into each well of the 384-well plates, followed by the addition of pNPP substrate (20 μL) and the absorbance at 405nm measured. Wells containing same percent of 0.1%DMSO served as vehicle control; Wells containing 1uM IL-17A-Inhibitor-1served as positive control.

The percent of inhibition of compounds were fitted with a 4-parameter logistic model using GraphPad Prism 8.0 to calculate the IC50 values according to the following formula: 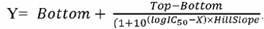

Y: %inhibition; X: log of inhibitor concentration; Bottom: %inhibition at 0; Top: %inhibition at 100; The Inhibition (%) for each compound calculated by equation:

Inhibition (%) = 100-(signalcmpd -signalAve_pc) / (signalAve_VC -signalAve_pc) ×100%.

SignalAve_pc: The average signal for the positive control across the plate.

SignalAve_vc: The average signal for vehicle control across the plate. signalcmpd: The average signal for the test compound.

### PAMPA Permeability Assay

Stock solutions (1 mM in DMSO) of test compounds and permeability controls (metoprolol, diclofenac, furosemide) were diluted to 10 μM in PBS (1% DMSO). Artificial membranes were prepared fresh by dissolving lecithin in dodecane (18 mg/mL). In the assay, 5 μL membrane solution was applied to filter plates, followed by 300 μL compound solution (donor) and 150 μL PBS/1% DMSO (acceptor). After 16 h incubation at 25°C (60 rpm), samples from both chambers were collected, protein-precipitated with cold acetonitrile (containing dexamethasone IS), and analyzed by LC-MS/MS. Effective permeability (Pe) was calculated using chamber volumes (donor: 0.3 mL; acceptor: 0.15 mL), incubation time (57600 s), and membrane area (0.24 cm^2^).

### Plasma Protein Binding Assay

Test compounds and warfarin (positive control) were prepared as 200 μM working solutions in DMSO, then diluted to 1 μM in EDTA-K2 plasma. Dialysis membranes (12-14 kDa) were pre-treated with water, 20% ethanol, and PBS buffer (pH 7.4). The assay was conducted in duplicate using 120 μL plasma samples dialyzed against equal volumes of PBS at 37°C (100 rpm) for 6 h. After incubation, plasma and buffer samples were protein-precipitated with cold acetonitrile (IS), centrifuged, and analyzed by LC-MS/MS. Percent free drug was calculated from buffer-to-plasma peak area ratios.

### Microsomal Stability Assay

Liver microsomes (20 mg/mL) were diluted to 0.5 mg/mL in phosphate buffer (pH 7.4). Test compounds and verapamil (positive control) were added at 1 μM final concentration. Reactions were initiated with NADPH (1 mM), and then aliquots were taken at 0, 5, 10, 15, 30, and 60 min, quenched with cold acetonitrile (IS), and centrifuged. Supernatants were analyzed by LC-MS/MS to determine remaining parent compound. In vitro half-life (t1/2) and intrinsic clearance (CLint) were calculated from logarithmic plots of concentration versus time.

### CYP450 Inhibition Assay

Human liver microsomes (0.1 mg/mL) were pre-incubated with test compounds and CYP-specific substrates: phenacetin (1A2), bupropion (2B6), amodiaquine (2C8), diclofenac (2C9), (S)-mephenytoin (2C19), dextromethorphan (2D6), and testosterone (3A4). Reactions were initiated with NADPH (1 mM) and stopped at 10 min (30 min for CYP2C19) by acetonitrile precipitation. Metabolite formation was quantified by LC-MS/MS. Inhibition percentage was calculated relative to control reactions without inhibitor. Selective inhibitors (e.g., ketoconazole for CYP3A4) were included as controls.

### Pharmacokinetic Study

Mice were treated with 3 mg/kg(iv), 3, 10 and 30 mg/kg (p.o.). Rats were treated with 2 mg/kg(iv), 6.25, 12.5, 25, 40, 50 mg/kg (p.o.). Monkeys were treated with 1 mg/kg(iv), 3, 10, 30 mg/kg(p.o.). For each mouse in all group, blood samples were collected at the time point of 0.25, 0.5, 1, 2, 4, 8 and 24 h post-dose. For each rat in all group, blood samples were collected at the time point of 0.25, 0.5, 1, 2, 4, 8, 12 and 24 h post-dose. For each dog in all group, blood samples were collected at the time point of 0.25, 0.5, 1, 2, 4, 8, 12 and 28 h post-dose Blood samples were sampled into EDTA-coated tubes (Eppendorf) and placed on ice until centrifugation to obtain plasma samples. The plasma samples were stored at -80°C until analysis. The concentration of compound in plasma samples was determined using a LC-MS/MS method. PK parameters were estimated by non-compartmental model using WinNonlin. **Establishment of Collagen-Induced Arthritis (CIA) model and dosing**

Rat collagen induced arthritis was established to evaluate the efficacy of the compounds. Male Wistar rats, aged 6 weeks, were used in the CIA efficacy experiment. The materials and reagents of the CIA efficacy experiment are listed in the following **Table 1**:

**Table 1.**

| **Materials and Reagents** | **Vendor** | **Cat#** |
| --- | --- | --- |
| Type II collagen | Chondrex | CDX20022 |
| Complete Freund's adjuvant | Sigma | F5881 |
| Micro-sized glass syringe | HAMILTON | 81220 |
| Pipette | Eppendorf | 3124000121 |
| 1.25% tribromoethanol solution | Nanjing Aibei Biotechnology | M2960 |

Type II collagen (CII) was emulsified 1: 1 with Complete Freund’s Adjuvant (CFA) on ice until a thick, stable emulsion was obtained (drop test in water). The emulsified preparation should be kept on crushed ice and injected within 1 hour. On Day 0 (primary immunization), emulsified CII in CFA is injected intradermally (i.d.) into the tail skin approximately 1.5 cm distal to the base of the tail at a volume of 200 µL per rat. On Day 7, a booster immunization was administered following the same protocol, but with a reduced volume of 100 µL of freshly prepared emulsion. After the CIA rat successful modeling, the rats were randomized into seven groups (n=6-8 per group) based on body weight stratification: CIA model group (vehicle, BID), **6** (LY3509754, 6mg/kg, BID), **8** (LEO, 25 mg/kg, BID), **9** (this study,6.25?12.5 and 25 mg/kg, BID) and **10** (DC-806,25 mg/kg, BID). At the same time, normal wild-type rats receiving the vehicle was served as the negative control in this study. Following the initiation of treatment, the arthritis scores were determined every 2-3 days on the extremities of each rat and the score for each rat is the sum of the limb scores (range, 0-16 points, with a maximum score in each rat of 16). Additionally, the thickness of the hind paws was measured to assess the inhibitory effects of different compounds on swelling.

## Reference

1. McGeachy, M. J.; Cua, D. J.; Gaffen, S. L., The IL-17 Family of Cytokines in Health and Disease. Immunity 2019, 50 (4), 892–906.

2. Amatya, N.; Garg, A. V.; Gaffen, S. L., IL-17 Signaling: The Yin and the Yang. Trends Immunol 2017, 38 (5), 310–322.

3. Gaffen, S. L.; Jain, R.; Garg, A. V.; Cua, D. J., The IL-23-IL-17 immune axis: from mechanisms to therapeutic testing. Nat Rev Immunol 2014, 14 (9), 585–600.

4. Mills, K.H.G., IL-17 and IL-17-producing cells in protection versus pathology. Nat Rev Immunol 2023, 23 (1), 38-54.

5. Berg, S. H.; Balogh, E. A.; Ghamrawi, R. I.; Feldman, S. R., A review of secukinumab in psoriasis treatment. Immunotherapy 2021, 13 (3), 201–216.

6. Baraliakos, X.; Braun, J.; Deodhar, A.; Poddubnyy, D.; Kivitz, A.; Tahir, H.; Van den Bosch, F.; Delicha, E. M.; Talloczy, Z.; Fierlinger, A., Long-term efficacy and safety of secukinumab 150 mg in ankylosing spondylitis: 5-year results from the phase III MEASURE 1 extension study. RMD Open 2019, 5 (2), e001005.

7. Liu, L.; Lu, J.; Allan, B. W.; Tang, Y.; Tetreault, J.; Chow, C. K.; Barmettler, B.; Nelson, J.; Bina, H.; Huang, L.; Wroblewski, V. J.; Kikly, K., Generation and characterization of ixekizumab, a humanized monoclonal antibody that neutralizes interleukin-17A. J Inflamm Res 2016, 9, 39–50.

8. Foulkes, A. C.; Warren, R. B., Brodalumab in psoriasis: evidence to date and clinical potential. Drugs Context 2019, 8, 212570.

9. Zhang, B.; Domling, A., Small molecule modulators of IL-17A/IL-17RA: a patent review (2013-2021). Expert Opin Ther Pat 2022, 32 (11), 1161–1173.

10. Velcicky, J.; Bauer, M. R.; Schlapbach, A.; Lapointe, G.; Meyer, A.; Vogtle, M.; Blum, E.; Ngo, E.; Rolando, C.; Nimsgern, P.; Teixeira-Fouchard, S.; Lehmann, H.; Furet, P.; Berst, F.; Schumann, J.; Stringer, R.; Larger, P.; Schmid, C.; Prendergast, C. T.; Riek, S.; Schmutz, P.; Lehmann, S.; Berghausen, J.; Scheufler, C.; Rondeau, J. M.; Burkhart, C.; Knoepfel, T.; Gommermann, N., Discovery and In Vivo Exploration of 1,3,4-Oxadiazole and alpha-Fluoroacrylate Containing IL-17 Inhibitors. J Med Chem 2024, 67 (18), 16692–16711.

11. Datta-Mannan, A.; Regev, A.; Coutant, D. E.; Dropsey, A. J.; Foster, J.; Jones, S.; Poorbaugh, J.; Schmitz, C.; Wang, E.; Woodman, M. E., Safety, Tolerability, and Pharmacokinetics of an Oral Small Molecule Inhibitor of IL-17A (LY3509754): A Phase I Randomized Placebo-Controlled Study. Clin Pharmacol Ther 2024, 115 (5), 1152–1161.

